# Maternal mRNA deadenylation is defective in *in vitro* matured mouse and human oocytes

**DOI:** 10.1101/2021.08.29.458073

**Authors:** Yusheng Liu, Wenrong Tao, Yiwei Zhang, Hu Nie, Zhenzhen Hou, Jingye Zhang, Zhen Yang, Jiaqiang Wang, Falong Lu, Keliang Wu

## Abstract

Oocyte *in vitro* maturation is a technique of assisted reproductive technology that was first introduced in patients with polycystic ovarian syndrome and is now used in most fertility clinics. Thousands of genes show abnormally high expression in *in vitro* maturated metaphase II (*in vitro* MII) oocytes compared with *in vivo* maturated metaphase II (*in vivo* MII) oocytes in bovines, mice, and humans^1-3^. However, the underlying mechanisms of this abnormal expression are still poorly understood. In this study, we use PAIso-seq1 to reveal a transcriptome-wide expression profile of full-length transcripts containing entire poly(A) tails in *in vivo* and *in vitro* matured mouse and human oocytes. Our results indicate that more genes are up-regulated than down-regulated in *in vitro* MII oocytes in both mice and humans. Furthermore, we demonstrate that the observed increase in maternal mRNA abundance is caused by impaired deadenylation in *in vitro* MII oocytes in both mice and humans. We also found that the cytoplasmic polyadenylation of dormant *Btg4* and *Cnot7* mRNAs, which encode key components of deadenylation machinery, is impaired in *in vitro* MII oocytes in mice and humans respectively, likely contributing to reduced translation and impaired global maternal mRNA deadenylation. Our findings highlight that impaired maternal mRNA deadenylation is a definite molecular defect in *in vitro* MII oocytes in both mice and humans. The findings here offer a new criterion for evaluating the quality of *in vitro* MII oocytes and a potential direction for improving *in vitro* maturation by fixing the dysregulated maternal mRNA deadenylation.

## Introduction

Female infertility is becoming an exacerbating reproductive problem, for which the assisted reproductive technology (ART) is an effective treatment^4^. *In vitro* maturated metaphase II (*in vitro* MII) oocytes were first introduced as an assisted reproductive technology for patients with polycystic ovarian syndrome (PCOS)^5^ and patients with severe ovarian hyperstimulation syndrome (OHSS) during previous *in vitro* fertilization (IVF) treatments^6^. Currently, *in vitro* MII oocytes can be adapted as an option in almost all areas of fertility clinics, including PCO-like ovaries, resistant ovary syndrome, previous failed IVF attempts, and oocyte maturation problems^7^. Additionally, *in vitro* MII oocytes can be useful for preserving fertility in situations such as emergency oocyte retrieval due to malignancies^8^. The first baby from donor *in vitro* MII oocytes was born in 1991^9^, and the first baby from the mother’s own *in vitro* MII oocytes was born in 1994^10^. As such, in recent years, *in vitro* MII oocytes have gained increasing attention for their feasibility, safety, reproducibility, cost-effectiveness, and lack of OHSS risk^10,11^.

Previous studies have demonstrated that embryos derived from *in vitro* MII oocytes have a lower success rates of preimplantation development, pregnancy, and birth than those derived from mature *in vivo* MII (*in vivo* MII) oocytes^12^. The nuclear and cytoplasmic maturation in *in vitro* MII oocytes determines the performance of their oocytes, the quality of the embryo, and clinical outcomes^7,13^. The nuclear maturation can be easily evaluated with a microscope based on the first polar body extrusion. The cytoplasmic maturation defects during the *in vitro* maturation process include altered spindle positioning, mitochondrial membrane potential, the number of endoplasmic reticulum clusters, and the cortical actin cytoskeleton thickness in *in vitro* MII oocytes compared with *in vivo* MII oocytes^14,15^. Global gene expression profiling using human whole-genome arrays provides compelling evidence for the relative developmental incompetence of *in vitro* MII oocytes^1^, demonstrating that over 2,000 genes were expressed at 2-fold or greater levels in *in vitro* MII oocytes compared with *in vivo* MII oocytes. *In vitro* MII oocytes in mice and bovines were also demonstrated elevated expression patterns^2,3^. Nonetheless, the mechanisms underlying cytoplasmic maturation defects and upregulated gene expression profiles in *in vitro* MII oocytes remain largely unknown.

As transcription is silent, post-transcriptional regulation of maternal mRNA plays a dominant role in oocyte maturation, particularly deadenylation-dependent maternal mRNA decay. The CCR4-NOT deadenylase and its adaptor *Btg4* are critical regulators for this process^16-18^. In addition, *Cnot7* and *Cnot6l*, which encode catalytic subunit of the CCR4-NOT deadenylase, and *Btg4* are dormant maternal mRNAs that needs to be translationally activated through cytoplasmic polyadenylation during mouse oocyte maturation^16,17^. We demonstrated that maternal mRNAs are subjected to deadenylation-dependent decay during oocyte maturation in mice, rats, pigs, and humans^19-23^. Therefore, we hypothesize that the global gene upregulation observed in the *in vitro* MII oocytes is caused by impaired global deadenylation. To test this hypothesis, we used the PAIso-seq1 method, which is capable of measuring poly(A) tail inclusive full-length transcriptome from a single oocyte^24^, to measure the transcriptome-wide poly(A) tails in *in vivo* and *in vivo* MII oocytes in both mice and humans. By comparing the transcriptome-wide poly(A) tail length distribution of maternal mRNA in *in vitro* MII and *in vivo* MII oocytes, we found that the maternal mRNA decay mediated by poly(A) tail deadenylation was impaired in *in vitro* MII oocytes in both mice and humans. Furthermore, we found that cytoplasmic polyadenylation is critical for the translational activation of *Btg4* and *Cnot7* mRNAs, both of which encode factors critical for global maternal mRNA deadenylation and were impaired in *in vitro* MII oocytes in mice and humans. Therefore, our findings reveal that impaired maternal mRNA deadenylation is a critical defect in *in vitro* MII oocytes of both mice and humans, and provide the groundwork for improving the quality of *in vitro* MII oocytes.

## Results

### Abnormal maternal gene expression in *in vitro* MII oocytes in mice and humans

The deadenylation and cytoplasmic polyadenylation of the poly(A) tail of maternal mRNA both play essential roles during oocyte maturation^25,26^. Therefore, we performed PAIso-seq1 to analyze the poly(A) tail inclusive transcriptome in *in vitro* and *in vivo* MII oocytes as well as the germinal vesicle (GV) oocytes in both mice and humans (Fig. 1a, f). We found that the overall transcriptional level of individual genes significantly decreased during maturation in both mice and humans (Fig. 1b, g). This is consistent with the well-known global reduction of maternal mRNA during mammalian oocyte maturation, as measured by RNA-seq or microarray^1,16,17^. The transcriptional level of individual genes was significantly higher in *in vitro* than in *in vivo* MII oocytes in mice and humans (Fig. 1b, g), indicating that global maternal mRNA decay is impaired in *in vitro* MII oocytes. We found that 4,789 genes increased, while only 343 genes decreased in *in vitro* compared to *in vivo* MII oocytes in mice (Fig. 1c), and 3,072 genes increased while only 1,352 genes decreased in *in vitro* compared to *in vivo* MII oocytes in humans (Fig. 1h). The up-regulated genes enrich in cellular metabolic process, organelle organization, cell cycle, as well as DNA repair in both mice and humans (Fig. 1d, i). This indicates the presence of conserved defects in *in vitro* MII oocytes between mice and humans. *Bmp15* is known to inhibit follicle maturation and is downregulated during this process^27^; we found that it is upregulated in *in vitro* compared to *in vivo* MII oocytes in both mice and humans (Fig. 1e, j). Our findings are consistent with previous global gene expression profiling comparing *in vitro* and *in vivo* MII oocytes in humans and mice^1,2^ and were similar when comparing pluripotent stem cells (PSCs)-derived MII oocytes and *in vivo* MII oocytes of mouse^28^. These results indicate that the impaired maternal mRNA decay is one conserved defect in *in vitro* MII oocytes in both mice and humans.

**Fig. 1.**
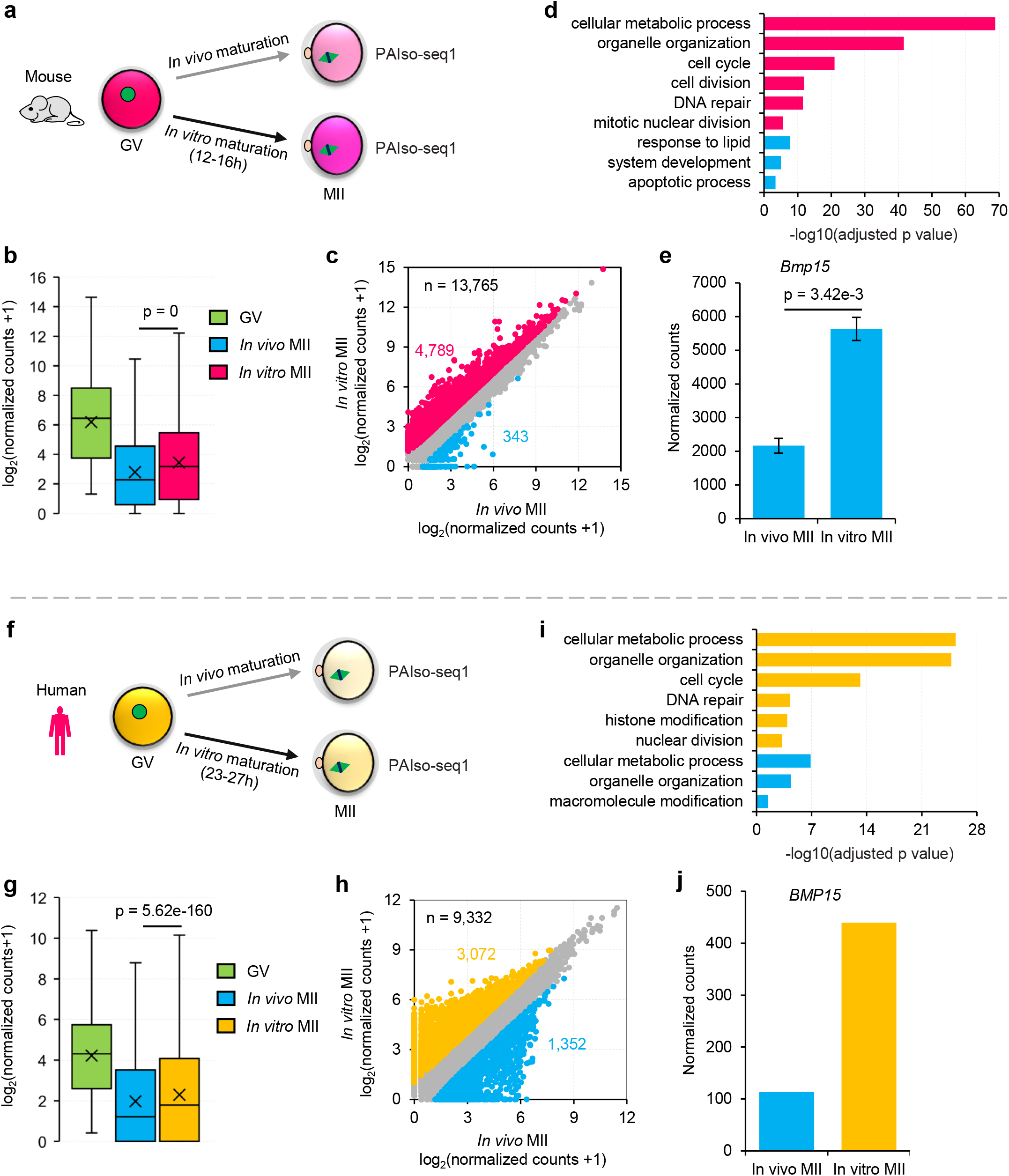
Abnormal maternal gene expression in mouse and human oocytes matured *in vitro*. **a, f**, Illustration of the *in vitro* and *in vivo* oocyte maturation experiments in mouse (**a**) and human (**f**) oocytes. **b, g**, Box plots for the normalized counts (in log2 scale) of genes (**b**, n = 14,131; **g**, n = 12,013) in germ-vesicle (GV) and *in vivo* maturated metaphase II (*in vivo* MII) or *in vitro* maturated metaphase II (*in vitro* MII) oocytes in mice (**b**) or humans (**g**). **c, h**, Scatter plot for the normalized counts of individual genes in MII oocytes matured *in vivo* or *in vitro* in mice (**c**) or humans (**h**). Each dot represents one gene. Genes with at least 1 read in one of the samples are included in the analysis. The number of genes included in the analysis and the numbers of differentially expressed genes are shown on the graphs. Genes upregulated in the *in vitro* MII oocytes are in red (**c**) or orange (**h**), while those downregulated in the *in vitro* MII oocytes are in blue. The differential expression is defined by a 2-fold cutoff. **d, i**, Gene Ontology (GO) analysis of genes upregulated (red in **d** and orange in **i**) or downregulated (blue) in the *in vitro* MII oocytes compared to *in vivo* MII oocytes in mice (**d**) or humans (**i**). **e, j**, Normalized counts of *Bmp15* in the *in vitro* and *in vivo* MII oocytes in mice (**e**) or humans (**j**). Error bars indicate the standard error of the mean (SEM) from two replicates. All *p-*values are calculated by Student’s *t-*test. For all the box plots, the “×” indicates the mean value, the black horizontal bars show the median value, and the top and bottom of the box represent the value of 25^th^ and 75^th^ percentile, respectively. The read counts are normalized by the counts of reads mapped to protein-coding genes in the mitochondria genome, if normalization is indicated.

### Impaired maternal mRNA deadenylation in *in vitro* MII oocytes of mouse and human

In eukaryotes, most mRNA decay is initiated by poly(A) tail deadenylation, including maternal mRNA decay during mammalian oocyte maturation^29^. It has been reported that deadenylation inhibition via *Btg4* depletion in mouse oocytes^16,17^ or *Btg4* mutation in human oocytes^18^ impedes maternal mRNA decay. Therefore, we explored whether the impaired maternal mRNA decay seen in *in vitro* MII oocytes is due to defective RNA deadenylation. To test this, we analyzed the transcriptome-wide poly(A) tail length distribution in *in vitro* and *in vivo* MII oocytes and found a global accumulation of transcripts with 20-100 nt poly(A) tails in *in vitro* MII oocytes in mice and humans (Fig. 2a, e). This indicates impaired maternal mRNA deadenylation, which is typically more severe in transcripts from genes that are upregulated in *in vitro* MII oocytes (Figs 1c, h, 2b, f), while transcripts from genes that are downregulated in *in vitro* MII oocytes showed an opposite pattern (Figs 1c, h, 2c, g). For example, *Bmp15* transcripts with 20-100 nt poly(A) tails were significantly accumulated in *in vitro* MII oocytes of both mice and humans (Fig. 2d, h). These results demonstrate that the deadenylation of maternal mRNA is impaired in *in vitro* MII oocytes, which is conserved in both mice and humans.

**Fig. 2.**
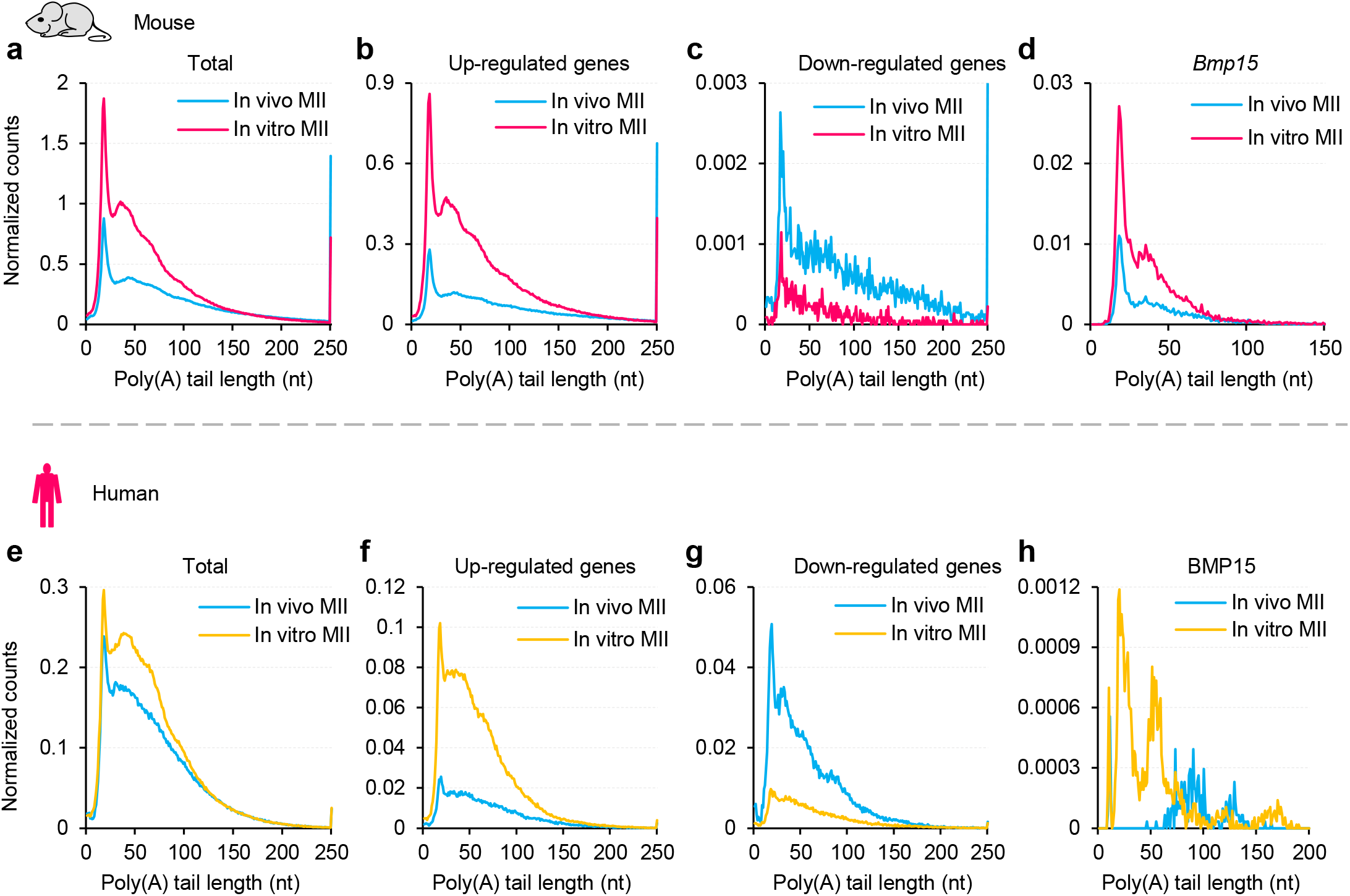
mRNA deadenylation is impaired in mouse and human oocytes matured *in vitro*. Histogram of poly(A) tails length of all transcripts (**a, e**), transcripts of the *in vitro* MII oocyte-upregulated genes (**b**, red genes (n = 4,789) in **Fig. 1c**; **f**, orange genes (n = 3,072) in **Fig. 1h**), transcripts of the *in vitro* MII oocyte-downregulated genes (**c**, blue genes (n = 343) in **Fig. 1c**; **g**, blue genes (n = 1,352) in **Fig. 1h**), or transcripts of *Bmp15* in *in vivo* or *in vitro* MII oocytes in mice (**a-d**) or humans (**e-h**). Histograms (bin size = 1 nt) are normalized by counts of reads mapped to protein-coding genes in the mitochondria genome. Transcripts with poly(A) tails of at least 1 nt are included in the analysis. Transcripts with poly(A) tail lengths greater than 250 nt (150 nt for **d** and 200 nt for **h**) are included in the 250 nt (150 nt for **d** and 200 nt for **h**) bin.

We report that non-A residues can be incorporated into maternal mRNA poly(A) tails through cytoplasmic polyadenylation followed by deadenylation during oocyte maturation, which is conversed in mice, rats, pigs, and humans^19-23^. When the deadenylation is impaired, the N (length between the end of 3′ UTR and the longest consecutive U, C, or G residues in poly(A) tails, Fig. 3a, c) is expected to be longer. This has been validated that the knockdown of *Btg4* in human zygotes and mouse MII oocytes results in longer N^20,21^. Therefore, we hypothesized that N should be longer in *in vitro* MII oocytes in which maternal mRNA deadenylation was impaired. As a result, we found that the N was longer for poly(A) tails with U residues in *in vitro* than in *in vivo* MII oocytes in both mice and humans (Fig. 3b, d). We also noticed that the ratio of poly(A) tails with C or G residues was lower in *in vitro* than in *in vivo* MII oocytes in mice and humans (Fig. 3b, d), reflecting lower incorporation of C and G residues (Extended Data Fig. 1). This suggests different mechanisms in U incorporation and C/G incorporation during oocyte maturation.

**Fig. 3.**
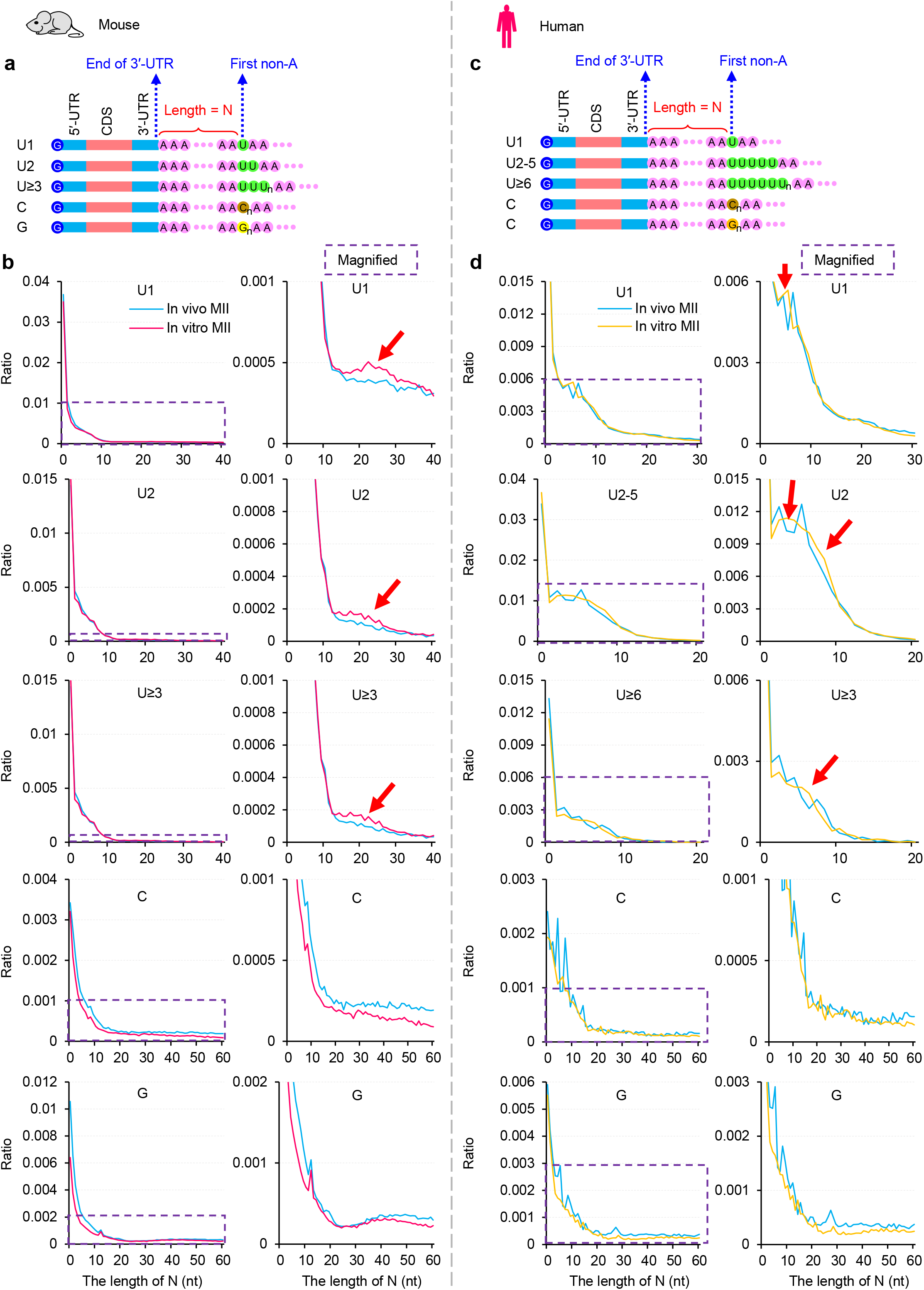
Mouse and human oocytes matured *in vitro* result in longer N. **a & c**, Diagram depicting mRNA with internal non-A residues in mice (**a**) and humans (**c**). N represents the length of residues between the end of 3′ UTR and the first base of the longest consecutive U, C, or G residues in a poly(A) tail. **b**, Histogram of the length of N and the ratio of U1, U2, U≥3, C, and G residues (from top to the bottom) in *in vivo* or *in vitro* MII oocytes in mice. **d**, Histogram of the length of N and the ratio of U1, U2-5, U≥6, C, and G residues (from top to the bottom) in *in vivo* or *in vitro* MII oocytes in humans. Histograms (bin size = 1 nt) are normalized to the total number of transcripts with a poly(A) tail of at least 1 nt. Magnified views of the regions in the purple dotted squares are shown on the right. Red arrows highlight the proportion of transcripts with longer N increases in *in vitro* MII oocytes.

Altogether, these data demonstrate that impaired maternal mRNA decay is caused by impaired deadenylation in *in vitro* MII oocytes in both mice and humans.

### Impaired cytoplasmic polyadenylation of *Btg4* and *Cnot7* in *in vitro* MII oocytes

Recent studies have demonstrated that CCR4-NOT deadenylase and its adaptor BTG4 are critical for global maternal mRNA deadenylation during mouse and human OET^16-18,21,22^. Therefore, we examined whether the impaired deadenylation in *in vitro* MII oocytes was caused by problems that occurred with CCR4-NOT deadenylase and its adaptor BTG4. The poly(A) tails of *Btg4* and *Cnot7* mRNA are short at the GV stage, and their translation is not initiated until cytoplasmic polyadenylation during oocyte maturation^16,17,30^. Therefore, we analyzed the transcript level and the poly(A) tail length of genes encoding components of the CCR4-NOT deadenylase and its adaptor BTG4 in *in vitro* MII oocytes of mice and humans.

At the transcript level, we found no or slight changes in the expression of these genes between *in vitro* and *in vivo* MII oocytes in mice and humans, except that *Cnot8* decreased dramatically in *in vitro* MII oocytes in humans (Fig. 4a, d). At the poly(A) tail length level, we observed a dramatic increase in poly(A) tail length for these genes during *in vivo* oocyte maturation in mice and humans (Fig. 4b, e), reflecting the cytoplasmic polyadenylation of these genes. However, we found that the cytoplasmic polyadenylation of *Btg4* in *in vitro* MII oocytes in mice and *CNOT7* in *in vitro* MII oocytes in humans was heavily impaired (Fig. 4b, e), suggesting that these factors cannot be translated at a normal level due to defective cytoplasmic polyadenylation.

**Fig. 4.**
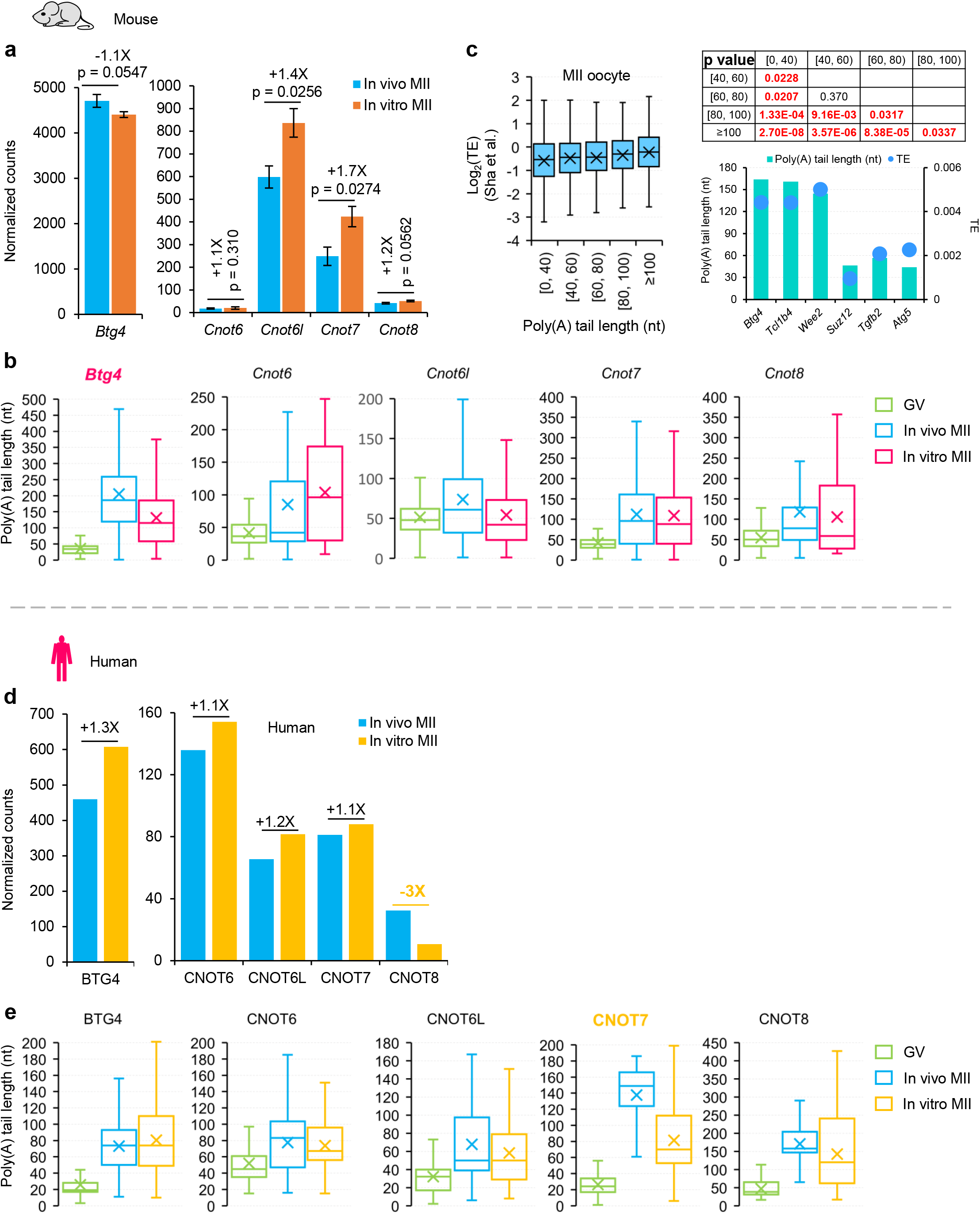
Polyadenylation of mRNAs encoding components of deadenylase complex is impaired in mouse and human oocytes matured *in vitro*. **a, d**, Normalized counts of *Btg4* (left) and the four *Cnot* family (right) genes (*Cnot6, Cnot6l, Cnot7, Cnot8*) in the *in vitro* and *in vivo* MII oocytes in mice (**a**) and humans (**d**). Error bars indicate the SEM from two replicates (n = 2). Fold changes for each gene are shown on top of the column. **b, e**, Box plot for the poly(A) tail length of *Btg4* and the four *Cnot* family genes in GV and MII oocytes matured *in vivo* or *in vitro* in mice (**b**) and humans (**e**). For all the box plots, the “×” indicates the mean value, the horizontal bars show the median value, and the top and bottom of the box represent the value of 25^th^ and 75^th^ percentile, respectively. **c**, Box plot of translational efficiency (TE, in log2 scale)^31^ of genes (n = 2,951) divided by the length of poly(A) tails in mouse *in vivo* MII oocytes. The *p*-values tested by Student’s *t*-test between each of the two groups are shown on the top right. Examples of poly(A) tail length and TE for *Btg4, Tcl1b4, Wee2, Suz12, Tgfb2*, and *Atg5* are shown on the bottom right.

The poly(A) tail length of mRNA is known to be positively associated with translational efficiency (TE) in mice GV to MI stage oocytes^22^. This positive association between poly(A) tail length and TE of mRNA has been suggested in mammalian MII oocytes but has never been examined transcriptome-widely. Therefore, we analyzed the relationship between our poly(A) tail length data and TE data from published polysome-seq in mouse MII oocytes^31^. We found that genes with longer poly(A) tails showed significantly higher TE in MII oocytes (Fig. 4c). For example, *Btg4, Tcl1b4*, and *Wee2* with poly(A) tail lengths around 140 nt showed much higher TE than *Suz12, Tgfb2*, and *Atg5* with poly(A) tail lengths around 50 nt (Fig. 4c). Altogether these results indicate that impaired deadenylation in *in vitro* MII oocytes is associated with the impaired cytoplasmic polyadenylation of mRNAs encoding CCR4-NOT deadenylase and its adaptor BTG4.

## Discussion

*In vitro* MII oocytes initially gained attention in ART for patients with PCOS^5^. More recently, it has been used in patients with repeated IVF failures^10^, resistant ovary syndrome, oocyte maturation problems^7^, and hormone-sensitive tumors^8^. Combining *in vitro* maturation and cryopreservation provides new opportunities for women to postpone motherhood^7^. The reduced ability of *in vitro* MII oocytes to support embryo development has been reported, however, the molecular mechanism underlying its defect is still unclear. In this study, we revealed that the maternal mRNA decay mediated by deadenylation is defective in *in vitro* MII oocytes in both mice and humans, which is associated with the defective cytoplasmic polyadenylation of mRNAs encoding CCR4-NOT deadenylase and its adaptor BTG4 (Fig. 5). These defects could potentially be used also as biomarkers in evaluating oocyte quality following the *in vitro* maturation procedure.

**Fig. 5.**
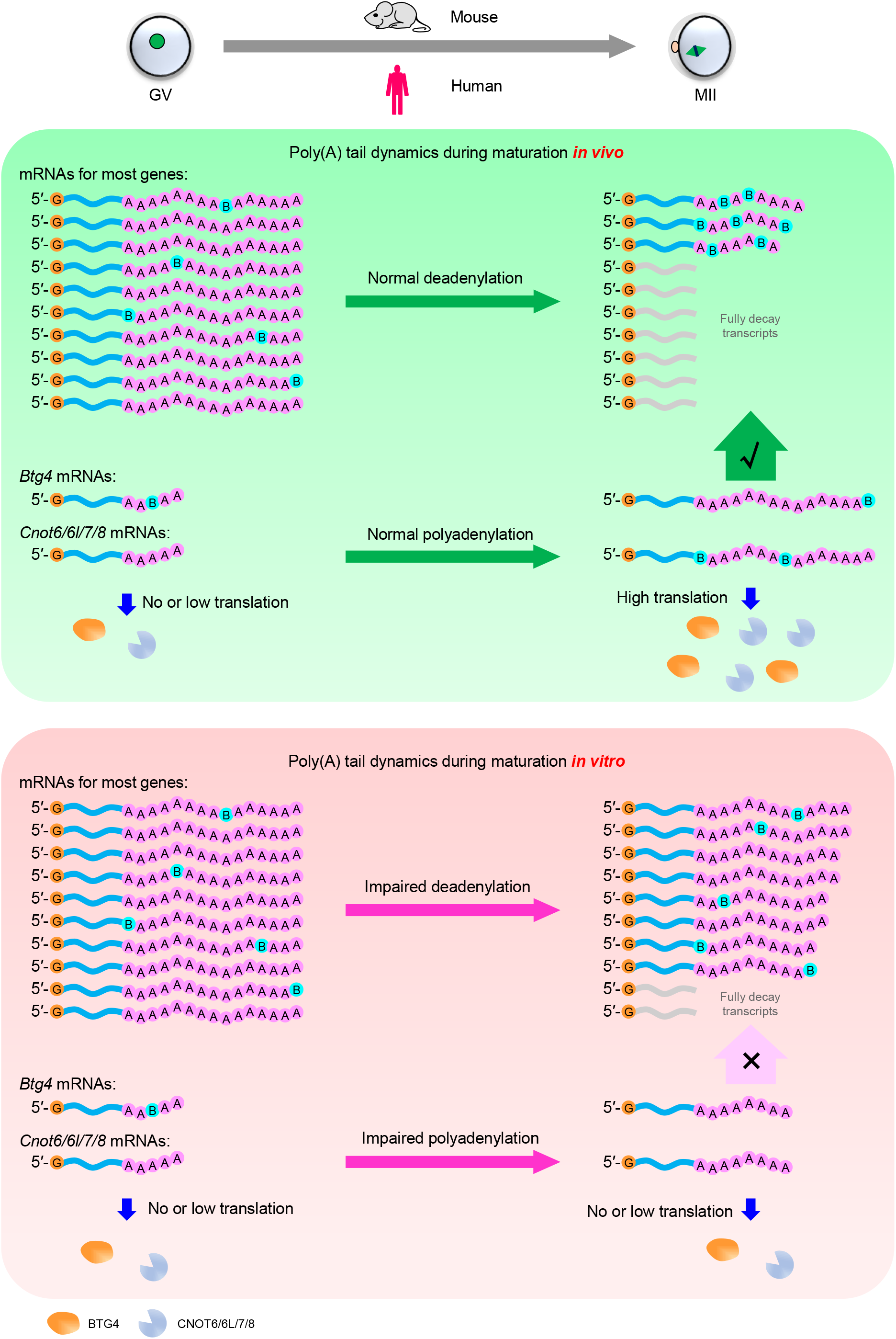
Summary of impaired maternal mRNA deadenylation during oocyte maturation *in vitro*. During oocyte maturation *in vivo* (up), the polyadenylation of mRNAs encoding components of deadenylase complex is normal, which results in high deadenylase abundance by high translation efficiency. Therefore, the global maternal mRNA deadenylation is normal. During oocyte maturation *in vitro* (down), the polyadenylation of mRNAs encoding components of deadenylase complex is impeded, which results in low deadenylase abundance by low translation efficiency. Therefore, the global maternal mRNA deadenylation is impaired.

In addition to impaired maternal mRNA decay, we revealed that the incorporation of non-A residues into poly(A) tails were also impaired during oocyte maturation, showing lower ratio of poly(A) tails with non-A residues, especially the C and G residues, in *in vitro* than in *in vivo* MII oocytes in both mice and humans (Extended Data Fig. 1). This was associated with the impaired cytoplasmic polyadenylation of *Tent4a/b* in *in vitro* MII oocytes which encode enzymes adding C and G residues to poly(A) tails^32^, although the expression levels of *Tent4a/b* were higher in *in vitro* than in *in vivo* MII oocytes of mouse and human (Extended Data Fig. 2). The mechanism behind how the cytoplasmic polyadenylation of *Btg4, Cnot7*, and *Tent4a/b* is impaired in *in vitro* MII oocytes warrants further exploration in the future.

BTG4-mediated maternal mRNA decay is essential for successful OET in mice and humans, since *Btg4*-null oocytes were infertile due to developmental arrest at the cleavage stage^16-18^. In this study, we demonstrated that maternal mRNA deadenylation is impaired in mouse and human *in vitro* MII oocytes, which is associated with decreased developmental potential for supporting embryonic development. Improving the activity of CCR4-NOT deadenylase and its adaptor BTG4 is likely to help solving this problem, and we expect that these approaches can one day promote the cytoplasmic maturation of oocytes under *in vitro* conditions. This will ultimately promote the future use of *in vitro* MII oocytes in human ART clinics.

Oocytes have recently been created *in vitro* from mouse pluripotent stem cells (PSCs), which can give rise to fertile and healthy offspring^28,33-36^. These groundbreaking results prove that it is possible to reproduce mice in the absence of natural oocytes. Using these methods in humans could provide hope for patients with infertility caused by oocyte abnormalities. Additionally, this procedure can contribute to regenerative medicine by providing oocytes for therapeutic cloning. Notably, PSC-derived MII oocytes also showed a higher level of gene expression of maternal mRNA than *in vivo* MII oocytes^28^. Therefore, our findings can also be used to evaluate the quality of and improve PSC-derived oocytes, a subject that warrants further study.

## Materials and Methods

### Human oocytes

The use of human gametes for this research follows the Human Biomedical Research Ethics Guidelines (set by National Health Commission of the People’s Republic of China on 1 December 2016), the 2016 Guidelines for Stem Cell Research and Clinical Translation (issued by the International Society for Stem Cell Research, ISSCR) and the Human Embryonic Stem Cell Research Ethics Guidelines (set by China National Center for Biotechnology Development on 24 December 2003). All the human related experiments in this study are in compliance with these relevant ethical regulations. The aim and protocols of this study has been reviewed and approved by the Institutional Review Board of Reproductive Medicine, Shandong University.

The donor women are 25–38 years old with tubal-factor infertility and their partners have healthy semen. Written informed consent was obtained from all oocyte and sperm donors. Oocytes donated from patients taking in-vitro fertilization treatments. Immature human GV oocytes without cumulus cells were matured in *in vitro* maturation (IVM) medium at 37 ° in an atmosphere with 5% CO_2_ for 24 hours. The IVM medium consists of M199 medium (GIBCO, 11-150-059) with 20% Systemic Serum Substitute (Irvine Scientific, 99193) and 75 mIU/mL of recombinant follicle stimulating hormone (Merck Serono). The oocytes were randomly assigned to experimental groups. Single oocytes were used for PAIso-seq1 analysis with 3 – 4 replicates for each stage.

### Mouse oocytes

CD1 (ICR) Mice were purchased from Beijing Vital River Laboratory Animal Technology Co., Ltd. and maintained according to the guidelines of the Animal Care and Use Committee of the Institute of Genetics and Development Biology, Chinese Academy of Sciences. GV oocytes were isolated from ovaries after injection with 10 U of pregnant mare serum gonadotropin (PMSG) (Prospec). GV oocytes without cumulus cells were *in vitro* cultured in M16 medium for 14 hours to collect *in vitro* matured MII oocytes. To obtain MII oocytes, the mice were injected with 10 U of PMSG and 10 U of human chorionic gonadotropin (hCG) (Prospec) at 46- to 48-hour intervals. MII oocytes were isolated from the oviduct, without mating, 14 hours after hCG injection.

### PAIso-seq1 library construction

The PAIso-seq1 libraries for human single oocytes were constructed following the single-cell PAIso-seq1 protocol according to previously described methods^24^. The PAIso-seq1 libraries for mouse oocytes were constructed with total RNA prepared from mouse oocytes using TRIzol reagent (Life Technologies). The libraries sequenced using PacBio Sequel II instruments at Annoroad.

## Data availability

The ccs data in bam format from PAIso-seq1 experiments will be available at Genome Sequence Archive hosted by National Genomic Data Center. This study includes analysis of the following published data: Sha et al. (Gene Expression Omnibus database (GEO) accession no. GSE118564). Custom scripts used for data analysis will be available upon request.

## Acknowledgements

This work was supported by the National Key Research and Development Program of China (2018YFC1004000),the Strategic Priority Research Program of the Chinese Academy of Sciences (XDA24020203), National Natural Science Foundation of China (31970588, 32170606, 81871168), Natural Science Foundation of Heilongjiang province (YQ2020C003), the China Postdoctoral Science Foundation (2020M670516, 2020T130687), the State Key Laboratory of Molecular Developmental Biology, and the Fundamental Research Funds of Shandong University.

## Author Contributions

Yusheng Liu, Jiaqiang Wang and Falong Lu conceived the project and designed the study. Yusheng Liu, Yiwei Zhang, Hu Nie, Jiaqiang Wang and Falong Lu analyzed the sequencing data. Wenrong Tao, Zhenzhen Hou, Jingye Zhang, Zhen Yang and Keliang Wu collected human oocytes. Yusheng Liu and Jiaqiang Wang organized all figures. Yusheng Liu, Jiaqiang Wang and Falong Lu supervised the project. Yusheng Liu, Jiaqiang Wang and Falong Lu wrote the manuscript with the input from the other authors.

## Figure legends

**Extended Data Fig. 1.**
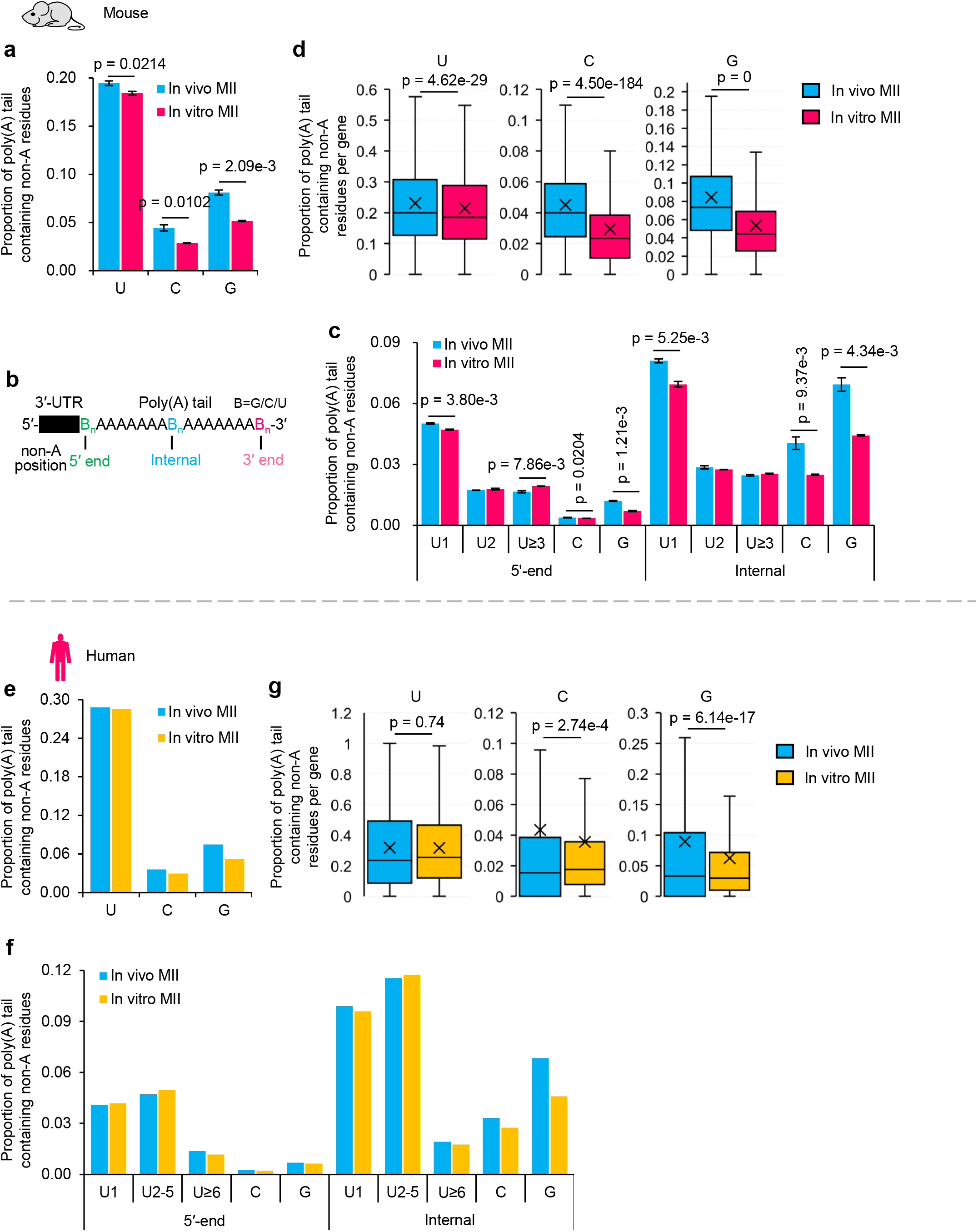
Proportion of transcripts containing non-A residues in the *in vitro* and *in vivo* MII oocytes in mice and humans. **a, e**, Overall proportion of poly(A) tails containing U, C, or G residues in the *in vitro* and *in vivo* MII oocytes in mice (**a**) and humans (**e**). Error bars indicate the SEM from two replicates and the *p*-value is calculated by Student’s *t*-test (**a**). **b**, Diagram depicting mRNA with 5′-end, internal, and 3′-end non-A residues. **c, f**, Proportion of poly(A) tails containing U, C, or G residues in 5′-end and internal in the *in vitro* and *in vivo* MII oocytes in mice (**c**) and humans (**f**). The U residues were further divided according to the length of the longest consecutive U (1, 2, and ≥3). **d, g**, Box plot for the proportion of reads containing U, C, or G residues of individual genes in the *in vitro* and *in vivo* MII oocytes in mice (**d**) and humans (**g**). Transcripts with poly(A) tails of at least 1 nt for a gene with at least 20 reads (**d**, n = 5,845; **g**, n = 3,163, 2,745, and 2,762 for U, C, and G, respectively) are included in the analysis. The *p*-value is calculated by Student’s *t*-test. For all the box plots, the “×” indicates the mean value, the black horizontal bars show the median value, and the top and bottom of the box represent the value of 25^th^ and 75^th^ percentile, respectively.

**Extended Data Fig. 2.**
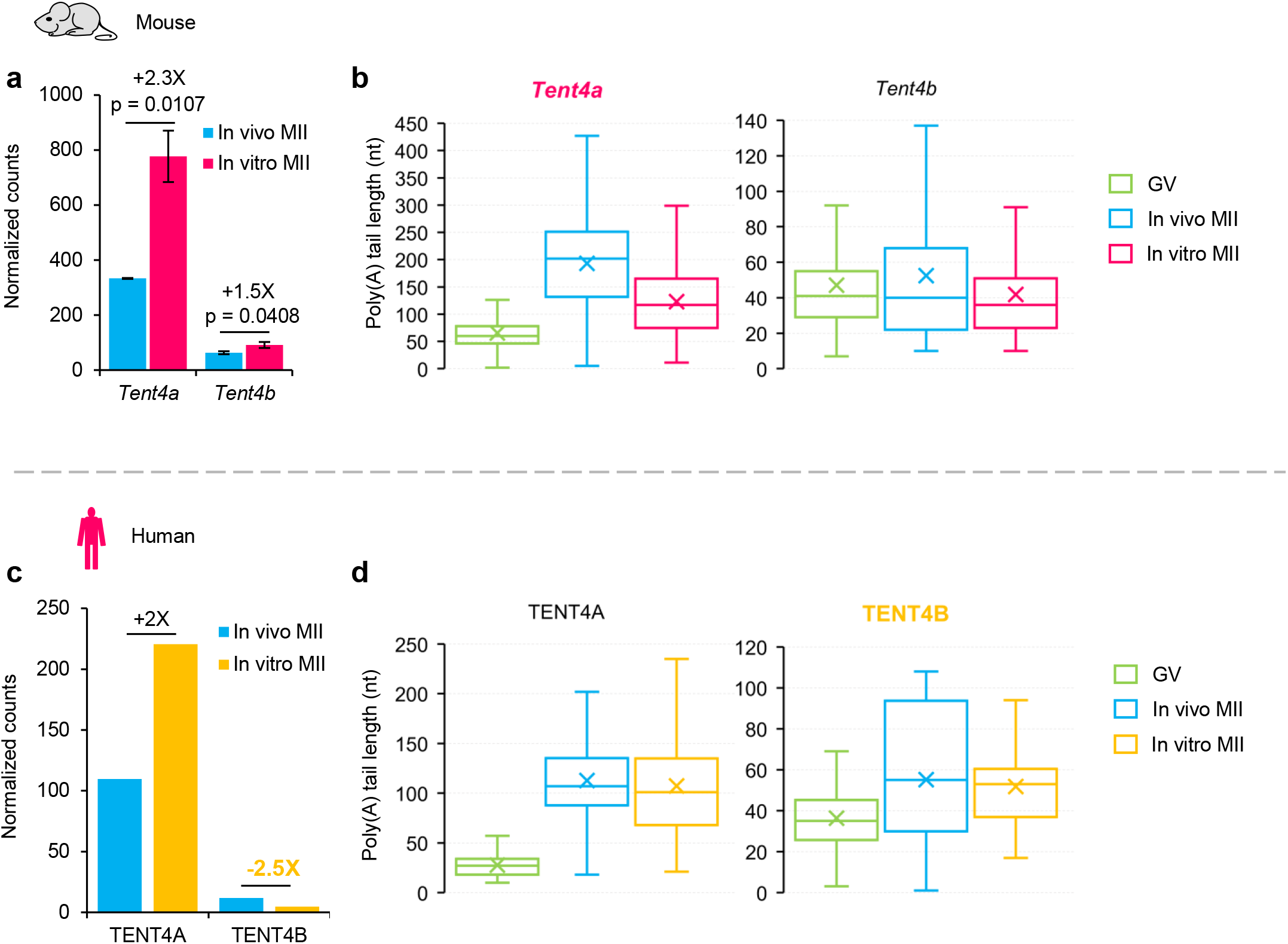
Expression and poly(A) tail length of *Tent4a/b* in the *in vitro* and *in vivo* MII oocytes in mice and humans. **a, c**, Normalized counts of *Tent4a* and *Tent4b* in the *in vitro* and *in vivo* MII oocytes in mice (**a**) and humans (**c**). Error bars indicate the SEM from two replicates and the *p*-value is calculated by Student’s *t*-test (**a**). **b, d**, Box plot for the poly(A) tail length of *Tent4a* and *Tent4b* in GV and MII oocytes matured *in vivo* or *in vitro* in mice (**b**) and humans (**d**). For all the box plots, the “×” indicates the mean value, the horizontal bars show the median value, and the top and bottom of the box represent the value of 25^th^ and 75^th^ percentile, respectively.

